# Differential activation of NLRP3 inflammasome by *Acinetobacter baumannii* strains

**DOI:** 10.1101/2022.04.29.490022

**Authors:** Fei-Ju Li, Lora Starrs, Anukriti Mathur, Hikari Ishii, Si Ming Man, Gaetan Burgio

**Affiliations:** Division of Immunity, Inflammation and Infection, The John Curtin School of Medical Research, the Australian National University, Canberra, ACT 2601, Australia; Division of Genome Sciences ad Cancer, The John Curtin School of Medical Research, the Australian National University, Canberra, ACT 2601, Australia

**Author notes:** Correspondence to: Gaetan Burgio, Division of Genome and Cancer, The John Curtin School of Medical, Research, The Australian National University, Canberra, ACT 2601, Australia.

**Keywords:** *Acinetobacter baumannii*, inflammasome, NLRP3, Caspase 11, Cell death

## Abstract

*Acinetobacter baumannii* is an emerging nosocomial, opportunistic pathogen with growing clinical significance globally. *A. baumannii* has an exceptional ability to rapidly develop drug resistance. It is frequently responsible for ventilator-associated pneumonia in clinical settings and inflammation resulting in severe sepsis. The inflammatory response is mediated by host pattern-recognition receptors and the inflammasomes. Inflammasome activation triggers inflammatory responses, including the secretion of the pro-inflammatory cytokines IL-1β and IL-18, the recruitment of innate immune effectors against *A. baumannii* infection, and the induction programmed cell death by pyroptosis. An important knowledge gap is how variation among clinical isolates affects the host’s innate response and activation of the inflammasome during *A. baumannii* infection. In this study, we compared nine *A. baumannii* strains, including clinical locally-acquired isolates, in their ability to induce activation of the inflammasome and programmed cell death pathway in primary macrophages and mice. We found a striking variability in survival outcomes of mice and bacterial dissemination in organs among three ATCC *A. baumannii* strains, likely due to the differences in virulence between strains. Interestingly, we found a stark contrast in activation of the NLRP3 inflammasome pathway, the non-canonical caspase-11 pathway, plasmatic secretion of the pro-inflammatory cytokines IL-1β and IL-18 between *A. baumannii* strains. Our study highlights the importance of utilising multiple bacterial strains and clinical isolates with differential virulence to investigate the innate immune response to *A. baumannii* infection.

## Introduction

*Acinetobacter baumannii* (*A. baumannii*) is a Gram-negative nosocomial pathogen commonly causing pneumonia and sepsis and known to have a high prevalence of harbouring multi-drug resistance (MDR) [1]. Due to its rapid development of antibiotic resistance, *A. baumannii* is one of the highly virulent ‘ESKAPE’ pathogens - a classification comprised of *A. baumannii, Enterococcus faecium, Staphylococcus aureus, Klebsiella pneumoniae, Pseudomonas aeruginosa,* and *Enterobacter* species [2]. Infections induced by these pathogens are responsible for life-threatening nosocomial infections in people who are most at risk, such as immunocompromised patients or those critically ill in the intensive care unit (ICU) [3].

Despite a rising incidence of multi-drug resistant *A. baumannii* infections, the immune mechanisms resulting from the infection remain elusive. The first line of host defence involves innate immune pattern-recognition receptors (PRRs) sensing conserved structures of microbial organisms, called pathogen-associated molecular patterns (PAMPs) [4]. The PRRs signalling cascades triggered upon bacterial recognition lead to TNFα and pro-IL-1β production, amplifying the inflammatory response. Pro-IL-1β is matured into the secretory form called IL-1β through the proteolytic activity of the cysteine protease Caspase-1 [5]. Mature IL-1β is an important pro-inflammatory cytokine, responsible for tightly regulating levels of inflammation in response to infection and activated by inflammasome sensor proteins, such as NLRP1, NLRP3 and NLRC4 [6, 7]. The role of the NLRP3 inflammasome has been characterized comprehensively for various diseases, including microbial infections. The assembly of the NLRP3 inflammasome – and thus the activation of Caspase-1 and secretion of IL-1β -following PRR signalling is termed the ‘canonical inflammasome pathway’ [8]. However, NLRP3 inflammasome activation may also be activated via a non-canonical inflammasome pathway involving Caspase-11 [8]. Caspase-11 is activated by direct binding to cytoplasmic Lipopolysaccharide (LPS); leading to Caspase-1-independent pyroptosis and NLRP3-Caspase-1-dependent release of IL-1β [4, 9].

In the limited studies on the immune response mounted against *A. baumannii*, there has been little investigation into the role of inflammasomes as host innate immune defence. It has previously been described that during *A. baumannii* infection, Toll-like receptor (TLR) 4 and other PRRs are responsible in propagating the signal for cytokine/chemokine production and inflammasome activation [10–13]. This pathway leads to the rapid and robust downstream effector responses. Depletion of neutrophils [14, 15] or macrophages [16] revealed an important role for these innate immune cells during *A. baumannii* infection. In addition, using an intra-nasal infection model, Kang and colleagues suggested a reduced lung pathology in NLRP3-deficient, Caspase-1/11-deficient and IL-1-receptor-deficient mice, although the bacteria burden, recruitment of immune cells and production of inflammatory cytokines and chemokines were not altered in these mice [17]. In contrast, Dikshit and colleagues showed that while the NLRP3 inflammasome contributes to host defence against *A. baumannii* clinical isolates, it is however not required to protect the host against a less virulent strain which is more sensitive to antibiotics [18].

Based on these findings, we hypothesised that inflammasome activation and programmed cell death are variable between the *A. baumannii* strains, correlating with bacterial virulence. To address this hypothesis, we examined the ability of three *A. baumannii* strains, frequently used in investigating bacterial biology, and six clinical isolates to induce pro-inflammatory cytokine production, inflammasome activation and programmed cell death. We indeed found a high variability in the survival of mice to infection with different strains of *A. baumannii* and variable bacterial dissemination in organs, likely due to the differences in strain virulence. Interestingly, we found a stark contrast between *A. baumannii* strains in the activation of the NLRP3 inflammasome pathways and secretion of pro-inflammatory cytokines by the host. This potentially correlated with the magnitude of programmed cell death observed and bacterial strain virulence. Holistically, our study underscores the importance of utilising multiple bacterial strains to investigate the host’s innate immune response to *A. baumannii* infection.

## Method

### Bacteria culture

*Acinetobacter baumannii* strains were originally purchased from American Type Culture Collection (ATCC). Strain *A. baumannii* Bouvet and Grimont ATCC 19606, *A. baumannii* type strain isolated from urine; strain *A. baumannii* Bouvet and Grimont (ATCC 17978), isolate from fatal meningitis of a four-months old infant; and strain *A. baumannii* ATCC BAA-1605 (short BAA-1605), a multi-drug resistant isolate from sputum of military personnel returning from Afghanistan entering a Canadian hospital. Additional clinical isolates including 834321, 820642, 916085, 938408, 834625 and 914394 were provided by the Canberra Hospital. Antibiotic resistance and isolation sites for each strain is listed in **Supplementary Table 1**. All bacteria strains were grown for 16-18 hours in Trypticase soy broth at 37 °C with shaking at 220 rotations per minute (rpm). For experiments herein, controls were set-up as ‘mock’ infection, that is, treated under identical conditions with the exception of adding bacteria.

### Cell culture

#### a. A549 cell line

The human carcinomic, lung epithelial line A549 was purchased from ATCC, and maintained in complete media consisting of Ham’s F-12K Medium (21127022, Gibco ThermoFisher) with 10% foetal bovine serum (FBS, F8192, Sigma) and supplemented with penicllin/streptomycin/glutamine (PSG, 10378016, Gibco ThermoFisher). Cells were maintained at 37C in 5% CO2 with >90% relative humidity. Cells were seeded in antibiotic-free media at a concentration of 1 × 10^5^ cells per well in 96-well plates the night prior to infection.

#### b. Bone Marrow Derived Macrophages (BMDMs)

Primary BMDMs were isolated from both mouse femurs and tibias under sterile conditions and in accordance with protocols mentioned in ‘Animals’. Briefly, bone marrow was flushed with RPMI with 2x penicillin/streptomycin and filtered through a 70um strainer (Cat No.:352350, BD Falcon). BMDM were isolated from red blood cells (RBC) through incubation of bone marrow with RBC lysis buffer (154.4mM NH_4_Cl, 9.99mM KHCO_3_ and 0.1267nM EDTA) and centrifugation at 1,500 rpm for 5minutes at room temperature. Cells were washed with RMPI with 2x penicillin/streptomycin and re-centrifuged at 1,500 rpm for 5 minutes at room temperature. Cells were seeded in a sterile 10-cm dish at a density of 5×10^6^ per dish in complete BMDM differentiation media consisting of RPMI-1640 media (22400-089, Gibco ThermoFisher) supplemented with 10% FBS, 1% penicillin and streptomycin (10378016, Gibco ThermoFisher) and 10ng/ml mouse GM-CSF (Cat. No.: 130-095-739, Miltenyi Biotech). Cultures were maintained at 37C in 5% CO2 with >90% relative humidity. Differentiation media was topped-up on day 3, and BMDMs were harvested on day 6 or 7 depending on confluency. BMDMs were harvested in 5uM EDTA/RPMI and centrifuged at 1,500 rpm for 5minutes at room temperature. seeded in antibiotic-free media at a concentration of 1 × 10^5^ cells per well in 96-well plates and incubated overnight at 37C in 5% CO2 with >90% relative humidity prior to infection.

### Quantification of cell death

#### a. Trypan blue

A549 cells were seeded into sterile glass bottom 24-well plate (Cat. No.: P24-1.5HN, Cellvis) at 4×10^5^ cells/well in antibiotic-free and serum-free media prior to *A. baumannii* inoculation. Cells were treated with different strains of *A. baumannii* at a multiplicity of infection (m.o.i.) of 80 for 24 hours (final volume 1 ml/well) at 37°C with 5% CO_2_ with >90% relative humidity. Cells were washed twice with sterile 1xPBS before staining with 2% trypan blue in 1xPBS at room temperature for 5-10 minutes. Post staining, cells were wash with sterile 1xPBS prior to fixation with 4% paraformaldehyde (PFA) (Cat. No.: 420801, BioLegend) at room temperature for at least one hour. PFA was carefully removed and 1xPBS was added to the wells for imaging. Cells were imaged using Zeiss Axio Observer under 40x objective. Five random fields were chosen per well and quantified using Image J with colour deconvolution plugin (version 1.64r).

#### b. Microscope

BMDMs were seeded into sterile glass bottom 24-well plate (Cat. No.: P24-1.5HN, Cellvis) at 4×10^5^ cells/well prior to *A. baumannii* inoculation. *A. baumannii* strains were prepared at m.o.i. of 10 and incubated with the cells for 24 hours (final volume 1 ml/well) at 37°C 5% CO_2_ and >90% relative humidity. Post infection, cells were washed twice with sterile 1xPBS before staining with Zombie Aqua (1:100, 100 μl/sample, Cat. No.: 423101, BioLegend) and Hoechst 33342 (1:50, 100 μl/sample, Cat. No.: 62249, Thermo Fisher Scientific) for 30 minutes in the dark at room temperature. Post staining, cells were washed twice and fixed in 4% PFA (Cat. No.: 420801, BioLegend) for 30 minutes in the dark, at room temperature. PFA was carefully removed and 1xPBS was added to the wells for imaging. Samples were imaged using Zeiss Axio Observer with an epifluorescence attachment and a digital camera. Five random fields were taken per well and quantified using Image J for mean staining area per channel (ver 1.64r).

### Animals

8-12-week-old mice were used for all mouse infection experiments and mice were between 8-16 weeks of age for BMDM isolation, C57BL/6J/^ANU^ mice were purchased from the Australian Phenomics Facility, the John Curtin School of Medical Research (Canberra, Australia) *Caspase-11^-/-^* mice were sourced from the Jackson Laboratory [19]. All mice were maintained under specific pathogen-free conditions. All animal studies were performed in accordance with the National Health and Medical Research Council code for the care and use of animals under the Protocol Numbers A2018-08 and A2021-14 approved by The Australian National University Animal Experimentation Ethics Committee.

### Mouse infection

#### a. Bacterial preparation and inoculation

C57BL/6J/^ANU^ mice were infected with 2×10^7^ CFU/mouse *A. baumannii* strains (ATCC 19606, ATCC 17978 or ATCC BAA-1605) intra-peritoneally. Briefly, single bacteria colony was grown for 16-18 hours in trypticase soy broth (TSB) at 37C, shaking at 220-250rpm. Next morning the cultures were refreshed in trypticase soy broth (1:5) and incubated at 37C with shaking for another 2h. The bacteria were then centrifuged at 2,300rpm for 30minutes and washed with sterile 1x PBS. The bacteria were resuspended in 200 μl PBS for a concentration of 2×10^7^ CFU/mouse. Actual inoculum concentrations were determined by plating serial dilutions on trypticase soy agar (TSA) plates.

#### b. Mouse monitoring and survival

Infected mice were monitored at 4, 8, 16, 20, and 28h post infection. Mice were monitored and scored based on condition, as assessed by factors including activity levels (slowing in movements etc.), grooming levels (smooth vs. rough coat), and hydration levels (drinking and eating). The endpoint of an experiment for each mouse was determined based on condition in accordance with the animal ethics protocol and the National Health and Medical Research Council code for the care and use of animals.

#### c. Quantification of bacterial burden

For enumeration of the bacteria burden in the lungs, liver, spleen and kidneys, mice were humanely euthanised between 8-20h post infections. and organs were aseptically removed prior to homogenisation in sterile 1xPBS. Approximately 20 to 50 mg of each organ were and filtered through a 70 μm nylon mesh cell strainer in 1 ml of sterile 1xPBS

For enumeration of bacterial burden in blood, blood was harvested in EDTA-containing tubes via post-euthanasia cardia-puncture. Serial dilutions of blood were plated on TSA plates to quantify bacterial load.

### Real-time reverse transcriptase polymerase chain reaction (real-time RT-PCR)

RNA was extracted from BMDMs using TRIzol (15596018, ThermoFisher Scientific). The isolated RNA was converted into cDNA using the High-Capacity cDNA Reverse Transcription Kit (4368813, ThermoFisher). RT-PCR was performed on an ABI StepOnePlus System PCR instrument with SYBR Green Real Time PCR Master Mixes (1725270, Bio-rad). Primer sequences can be found in Supplementary Table 2.

### Enzyme-linked immunosorbent assay (ELISA)

For cytokine measurement, plasma was collected after 8–20 h for analysis by ELISA. Serial diluted standard with known concentration was included in each plate. Cytokine concentrations from BMDMs were measured using TNFα (88-7324-77, Invitrogen), or IL-1β (88-7013-77, Invitrogen) ELISA kits according to the manufacturers’ instructions. All plates were measure using TECAN Infinite® 200 Pro (Tecan, Männedorf, Switzerland), with wavelength set at 450 nm.

### Western blotting

#### a. Sample preparation

Post infection, BMDMs and supernatant sample were lysed in Radioimmunoprecipitation assay buffer (RIPA) lysis buffer supplemented with protease inhibitors, i.e., Complete Protease Inhibitor Cocktail Tablets (Cat No.: 04693132001, Roche) to prevent sample degradation. Samples were heated at 95°C for 5 minutes with 6x Laemlii buffer containing Sodium Sodecyl Sulfate (SDS) and 100 mM Dithiothreitol (DTT) for 5 minutes before storing at −80°C.

#### b. Sodium Dodecyl Sulfate Polyacrylamide Gel Electrophoresis (SDS-PAGE)

Samples were thawed on ice and then denatured at 95°C for 10 minutes. Each sample was loaded on an individual lane of a 4-15% gradient SDS-PAGE gel (Cat No.: 456-1086, Bio-Rad) and run under reducing conditions with a constant voltage of 200 volts for approximately 25 minutes until the dye front reached the end of the gel. The resolved proteins in the SDS-PAGE gel were then transferred to a 0.45 μm Polyvinylidene fluoride (PVDF) membrane (Cat No.: 1620115, Bio-Rad) by electroblotting. An electric current of with 400 mA was applied to the apparatus for 1.5 hours at 4°C. Following the transfer, the membrane was blocked with 5% (w/v) skim milk in PBS for 1 hour at room temperature to prevent non-specific binding of Immunoglobulins (Ig).

#### c. Detection of protein of interest

Post-blocking the PVDF membrane was incubated with primary anti-caspase-1 (1:1000, Cat. No.: 106-42020, Adipogen), caspase-11 (1:1000, Cat. No.: NB120-10454, Novusbio), or Glyceraldehyde 3-phosphate dehydrogenase (GAPDH) (1:1000, Cat No.: MAB374, Merck Millipore), diluted in 1% (w/v) skim milk in PBST (1xPBS with 0.1% Tween-20) overnight at 4°C with gentle rocking. PVDF membranes were then incubated with species-specific horseradish peroxidase-conjugated secondary antibodies (1:5000) for 1 hour at room temperature with gentle rocking. Immunoreactive proteins were detected by applying ECL Western blotting Detection Reagent (Cat No.: 1705060, Bio-Rad) or SuperSignal™ West Femto Maximum Sensitivity Substrate (Cat. No.: 34096, Thermo Fisher Scientific).

### Statistical analysis

The GraphPad Prism 9.0 software was used for data analysis. Data are shown as mean ± s.e.m. Statistical significance was determined by t-tests (two-tailed) for two groups or one-way analysis of variance (with Dunnett’s multiple-comparisons test) for three or more groups. P < 0.05 was considered statistically significant.

## Result

### Multi-drug resistant *A. baumannii* ATCC BAA-1605 induced the highest lethality and bacterial burden

To assess the virulence of the *A. baumannii* strains ATCC 19606, 17978 and 1605, we examined tissue-specific bacterial burden and lethality in mice. Using the same dose (2×10^7^ CFU/mouse via intraperitoneal inoculation), the strain ATCC 17978 induced no lethality **(Fig. 1a)**, low bacterial load in blood **(Fig. 1b)**, and low colonisation level (< 100 CFU/mg) in the lung, liver, spleen, and kidney 16-20 hours post inoculation **(Fig. 1c)**. In contrast, the MDR strain ATCC BAA-1605 induced lethality in 62% of the mice **(Fig. 1a)**, with a high bacterial burden in the blood **(Fig. 1b)**, lung, liver, spleen or kidney **(Fig. 1c),** while the virulence and colonisation rat of the ATCC 19606 strain was intermediate between the ATCC 17978 and the ATCC BAA-1605 strains. Next, we compared the level of pro-inflammatory cytokines in the plasma of infected mice. Both the strain ATCC 19606 and the MDR strain ATCC BAA-1605 induced higher plasmatic IL-18 and IL-1β secretion level compared to the avirulent ATCC 17978. Together, these results suggest these *A. baumannii* strains differentially colonise, trigger inflammation and pro-inflammatory cytokine release, leading to differences in survival.

**Figure 1.**
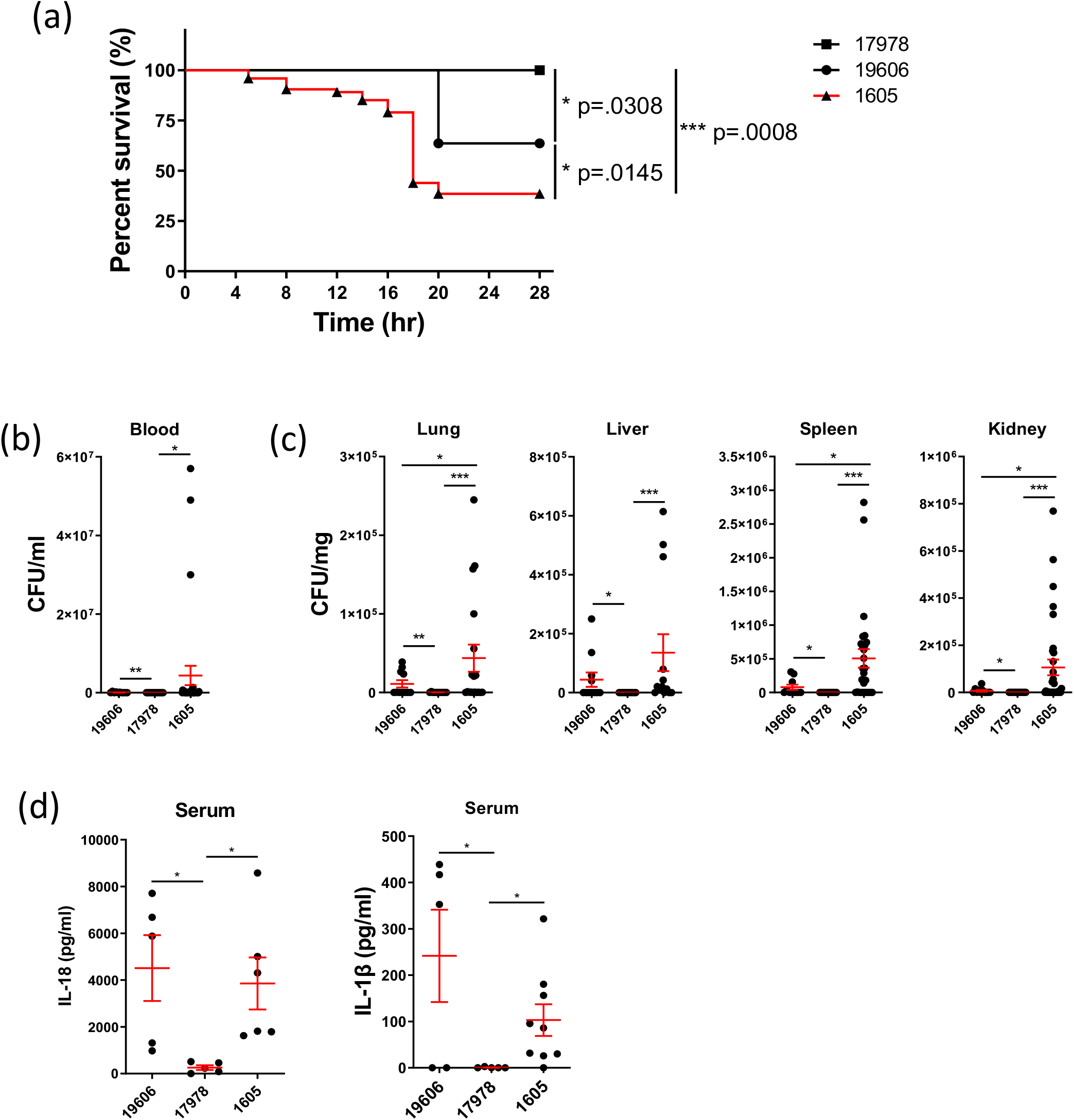
(a) The survival of wild-type mice following infection by three different strains of *A. baumannii* (i.p. 2×10^7^CFU/mouse). The level of (b) bacteraemia and (c) bacteria dissemination to different organs at 16-20 hours post infection was quantified. (d) Serum levels of IL-18 and IL-1β in canonical knockout mice post 16-20 hours intraperitoneal *A. baumannii* infection. Data were collected from at least three independent experiments, n ≥ 11. (b, c, d) data are shown as mean ± SEM. *, P < 0.05, **, P < 0.01, ***, P < 0.001.

### Differential upregulation of inflammasome components by different *A. baumannii* strains

We next determined which inflammasome is responsible for the maturation and release of pro-inflammatory cytokines IL-1β and IL-18. We checked the expression of inflammasome sensors via qRT-PCR on bone marrow derived macrophages (BMDMs) over for 24 hours of infection. For three *A. baumannii* strains, induction of NLRP3 and Caspase-11 was observed in BMDMs six hours post inoculation, which was sustained until 12 hours **(Fig. 2a)**. Interestingly, we observed a prolonged 8.4-fold induction of Caspase-11 24 hours post inoculation in BMDMs inoculated with the strain ATCC 17978 compared to ATCC 19606 or MDR strain ATCC BAA-1605. The MDR ATCC BAA-1605 strain induced a modest two-fold increase NLRC4 expression 24 hours post inoculation compared to mock (where a fold induction of 1 is equivalent to mock treatment) **(Fig. 2a)**. We noted no induction of the gene encoding AIM2 by any of the *A. baumannii* strains tested. We found no difference in the pro-inflammatory cytokines TNFα or IL-1β levels among the three strains from 6 to 24 hours post inoculation in BMDMs by qRT-PCR **(Fig. 2b)** or ELISA **(Suppl. Fig. 1)**. Overall, our findings suggest a consistent and preferential upregulation of NLRP3 and caspase-11 in response to several strains of *A. baumannii*.

**Figure 2.**
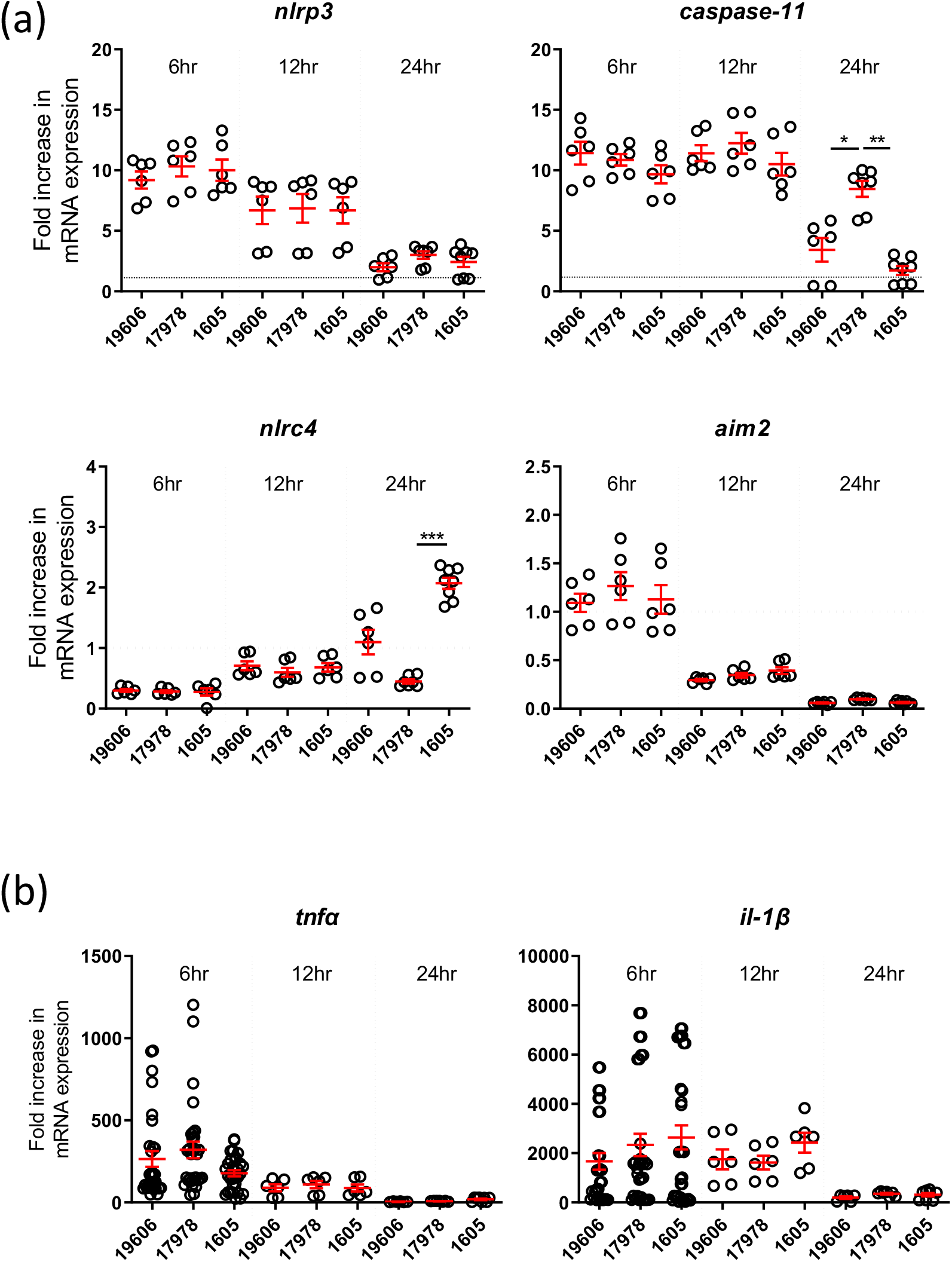
Mouse BMDMs were infected with m.o.i. 10 of live *A. baumannii* and analysed. (a) qRT-PCR transcript levels of *Nlrc4, Aim2, Nlrp3* and *Caspase-11*. (b) Tnfα and IL-1β, produced by wild-type mouse BMDMs between 6-24hours post-infection (lysate), n=6. mean ± SEM. *, P < 0.05, **, P < 0.01, ***, P < 0.001

We next hypothesised the magnitude and type of inflammatory response to *A. baumannii* is strain-dependent. BMDMs were infected with six *A. baumannii* clinical strains with various degree of resistance to antibiotics, including Carbapenem **(Suppl. Table 1)**. We firstly quantified Caspase-1, Caspase-11 and gasdermin D (GSDMD) cleavages by immunoblotting in WT BMDMs 12 hours post infection. We found the multidrug resistant strains 834321 and 916085 seem to be activating the highest level of Caspase-1, Caspase-11 and GSDMD-dependant cell death **(Fig. 3a)**. To confirm, we performed further analysis by measuring the associated cytokine IL-1β level in the supernatant of WT and *Caspase 11^-/-^* BMDMs 12 hours post infection. We noted a high variation in secreted IL-1β between all the clinical strains tested. Again, the multidrug resistant strains 834321 and 916085 seem to be inducing the highest levels of IL-1β secretion in WT BMDMs, but not in *Caspase 11^-/-^* BMDMs **(Fig. 3b)**. **(Fig. 3b)**. Together it suggests high variability in the inflammasome activation and IL-1β between and among the tested *A. baumannii* strains.

**Figure 3.**
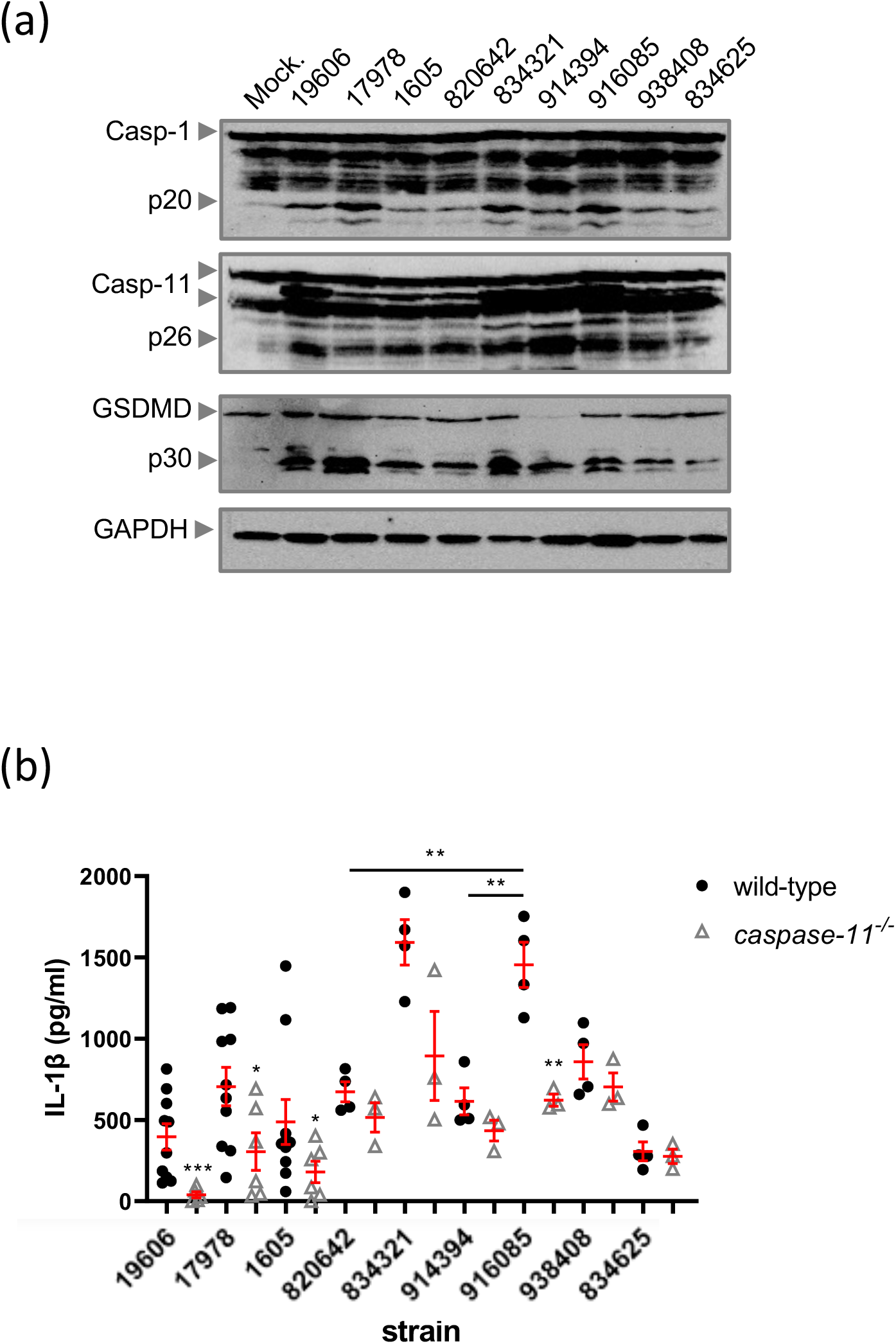
(a) Representative western blot image of caspase-1, caspase-11, and GSDMD post infection with different *A. baumannii* strains. (b) ELISA of IL-1β levels produced by mouse BMDM after infection m.o.i. 10 of *A. baumannii* infection for 12 hours. n≥3. mean ± SEM. *, P < 0.05, **, P < 0.01, ***, P < 0.001

### *A. baumannii* infection induces host programmed cell death

The activation of GSDMD and the release of IL-1β suggest that *A. baumannii* strains were likely to induce pyroptosis in BMDMs. We therefore quantified the level of cell death in BMDMs using the cell-permeability dye Zombie aqua, which detects cells with damaged membranes. We observed a variation in the level of cell death between BMDMs infected with different strains, with ATCC 17978 being the lowest **(Fig. 4a, b)**, which might be explained by its low virulence **(See Fig. 1)**. To investigate this further, we next determined whether this trend would be observed for another cell type other than BMDMs. As *A. baumannii* preferentially invades lung epithelium [20], we quantified cell death in the human A549 lung epithelial cell line. We infected A549 cells with *A. baumannii* and analysed using Zombie aqua and trypan blue uptake assays **(Suppl Fig. 2)**. Based on trypan blue and Zombie aqua staining, the strain ATCC BAA-1605 induced the highest level of cell death when compared to ATCC 17978 and ATCC 19606 strains **(Fig. 4a, b)**. We next postulated we would observe a high variation in cell death in response to infection by clinical strains. Amongst all strains tested, Trypan blue staining revealed that the clinical strain 916085 induced the highest level of cell death in A549 cells **(Suppl Fig. 2)**. Overall, it suggests a high variability of the *A. baumannii* strains in inducing cell death.

**Figure 4.**
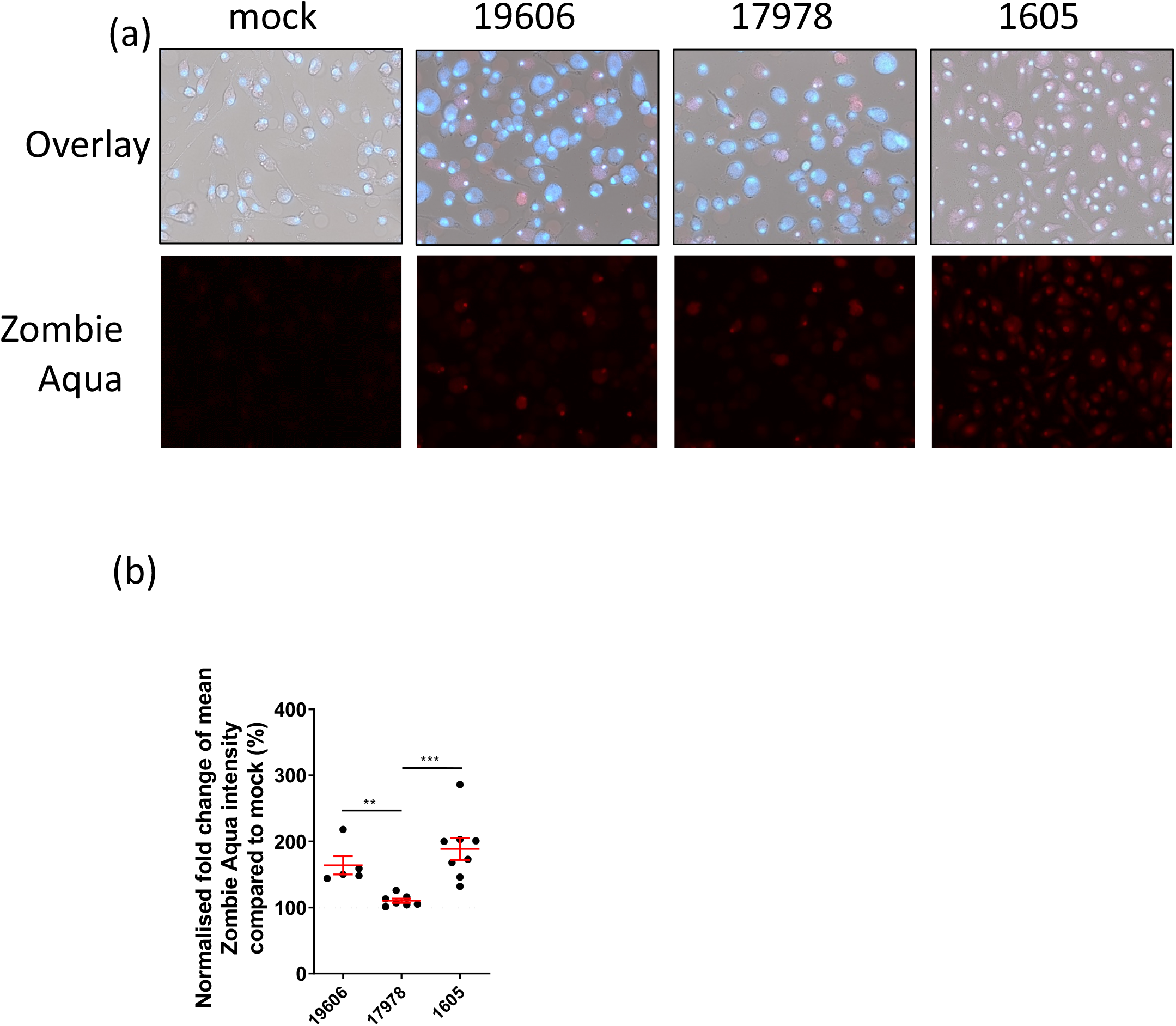
(a) representative image of wild-type mouse BMDM cell death post m.o.i. 10 of A. *baumannii* infection for 24 hours with Hoechst (blue) and zombie aqua (red) staining. Brightfield staining was included (top panel) (b) Densitometry quantification of the Hoechst and Zombie aqua staining from 5 random field per samples. N>3. All data are shown as mean ± SEM. *, P < 0.05, **, P < 0.01, ***, P < 0.001

## Discussion

*Acinetobacter baumannii* is a Gram-negative bacterium that causes opportunistic infections in humans. Despite its clinical significance, there is a lack of studies evaluating strain-dependency in the induction of inflammation and cell death responses. We examined strain-to-strain variation in virulence, pro-inflammatory cytokine secretion, inflammasome activation and programmed cell death. We observed a notable difference in virulence, pro-inflammatory cytokines, caspase-11 activation and programmed cell death in three commercially available *A. baumannii* strains ATCC 17978, 19606 and BAA-1605 commonly used to investigate *A. baumannii* pathogen biology and virulence. Here, we show that MDR ATCC BAA-1605 is more virulent than the ATCC 19606 and 17978 strains. Consistent with a previously published study [21], we found that *A. baumannii* is rapidly disseminated via the blood to peripheral organs, including the lung and spleen. Additionally, we show that the least virulent strain 17978 induced minimal systemic inflammatory responses, suggesting the acute lethality of other strains might be dependent on the degree of inflammatory response and pro-inflammatory cytokine releases.

Given the differential pro-inflammatory cytokine release and programmed cell death induced by different *A. baumannii* strains in mice, we wondered which inflammasome sensor and pathway was activated. Consistent with previous studies [17, 18, 22], we found an induction of NLRP3 and Caspase-11 inflammasome components early in the infection. Interestingly, neither the extended induction of Caspase-11 by the least virulent strain 17978 nor the delayed induction of NLRC4 by MDR ATCC BAA-1605 correlated with different levels of pro-inflammatory cytokines released in BMDMs. We confirmed this observation using six additional clinical isolates which exhibit various degrees of antibiotic resistance. We have titrated different m.o.i. versus IL-1β secretion and noted a lack of dose dependency (**Suppl Fig. 1**). We reasoned that the activation of inflammasomes largely depended on live *A. baumannii* activity rather than the presence of virulence factors.

We examined programmed cell death in BMDMs and a human epithelial cells using three commercial *A. baumannii* strains and six additional clinical isolates with various antibiotic resistance profiles. We found that the multidrug resistant clinical isolate 916085 induced the highest cell death. Interestingly, we have found a trend in relation to the variation in cell death versus inflammasome activation and pro-inflammatory cytokine secretion in our commercial and clinical strains. This suggests *A. baumannii* virulence factors are potentially modulating inflammasome activation and programmed cell death. Previous work demonstrated the role of virulence factors such as the IAV BP1-F2 [23], the UPEC alpha-hemolysin [24] and toxins of *Bacillus cereus* [25, 26] in determining NLRP3 activation and severe pathogenicity. Recently, the virulence-related OmpA was identified as an activator of NLRP3 inflammasome via Caspase-1 in *A. baumannii* infections [27]. We performed a multiple sequence alignment of the Omp38 protein sequence between the strains ATCC 17978, 19696 and BAA 1605 and found no differences in the OmpA domain (data not shown). Further analyses would assist with detailed characterisation of additional virulence factors in *A. baumannii* which may be responsible for Caspase-11 and NLRP3 activation leading to pyroptosis.

Altogether, we identified a correlation between *in vivo* virulence and *in vitro* host cell cytotoxicity for the three ATCC *A. baumannii* strains tested. We found that compared to ATCC type strain 19606 or strain 17978, MDR ATCC BAA-1605 was the most virulent strain both *in vitro* and *in vivo.* We identified that the induction of different inflammasome transcripts does not directly equate to the level of inflammation *in vitro*. Additionally, due toa lack of dosedependency *in vitro*, the activation of inflammasome was thought to be largely dependent on bacterial vitality. We propose that the correlation between *in vitro* cell death and *in vitro* Caspase-11-dependency may be harnessed as a screening tool to select for more virulent strains *in vivo*.

## Acknowledgement

We would like to thank Mr Mick Devoy and Dr Harpreet Vora, the Biomolecular resource facility and the Australian Phenomics Facility for technical assistance. The authors would like to acknowledge Dr Karina Kennedy for providing the clinical *A. baumannii* strains. F.J.L is supported from the Taiwan-Australian National University Scholarship. H.I is supported from the TOBITATE young ambassador program.

## Authors contribution

G.B., F.J.L. and L.S. conceptualised the study. F.J.L., L.S., A.M., H.I. performed the study. G.B., F.J.L., L.S., A.M., H.I. and S.M.M analysed the data. G.B. and F.J.L. wrote the manuscript. G.B. supervised the overall study. All authors reviewed and approved the manuscript.

---

**Table 1.** ATCC and clinical *A. baumannii* strain isolation sites and antibiotic resistance profiles.

| Strain | Origin | Isolation site | Resistant to |
| --- | --- | --- | --- |
| ATCC 19606 | ATCC | Urine | NA |
| ATCC 17978 | ATCC | Fatal meningitis | NA |
| ATCC BAA-1605 (1605) | ATCC | Sputum | Ceftazidime, Gentamicin, Ticarcillin, Piperacillin, Aztreonam, Cefepime, Ciprofloxacin, Imipenem, Meropenem |
| 916085 | Canberra Hospital | Swab | Ciprofloxacin, Gentamicin, Meropenem, Tobramycin |
| 834625 | Canberra Hospital | Blood | Ceftriaxone, Trimethoprim |
| 938408 | Canberra Hospital | Blood | Amp/Amoxicillin, Amox/Clav Acid, Cefazolin |
| 820642 | Canberra Hospital | Blood | Amp/Amoxicillin, Amox/Clav Acid, Cefazolin, Ceftriaxone, Trimethoprim |
| 834321 | Canberra Hospital | Urine | Ceftriaxone, Ciprofloxacin, Gentamicin, Norfloxacin, Tobramycin, Meropenem |
| 914394 | Canberra Hospital | Urine | Amikacin, Cefepime, Ceftazidime, Ciprofloxacin, Gentamicin, Meropenem, Pip/Tazobactam, Tica/Clax Acid, Tobramycin |

**Table 2.** Mouse primers used for qRT-PCR.

| Name | Forward (5'-3') | Reverse (5'-3') |
| --- | --- | --- |
| *Tnfa* | gcc tct tct cat tcc tgc ttg | ctg atg aga ggg agg cca tt |
| *Il1b* | gac ctt cca gga tga gga ca | agc tca tat ggg tcc gac ag |
| *Aim2* | gat tca aag tgc agg tgc gg | tct gag gct tag ctt gag gac |
| *Caspase-11* | aca atg ctg aac gca gtg ac | ctg gtt cct cca ttt cca ga |
| *Nlrp3* | gtg gtg acc ctc tgt gag gt | tct tcc tgg agc gct tct aa |
| *Nlrc4* | cta cat tga tgc tgc ctt gg | tct ctt cgt ctc tga gtc tc |
| *Gapdh* | gag gaa cct gcc aag tat g | tgg gag ttg ctg ttg aag |

## Supplementary material

**Supplementary Figure 1:**
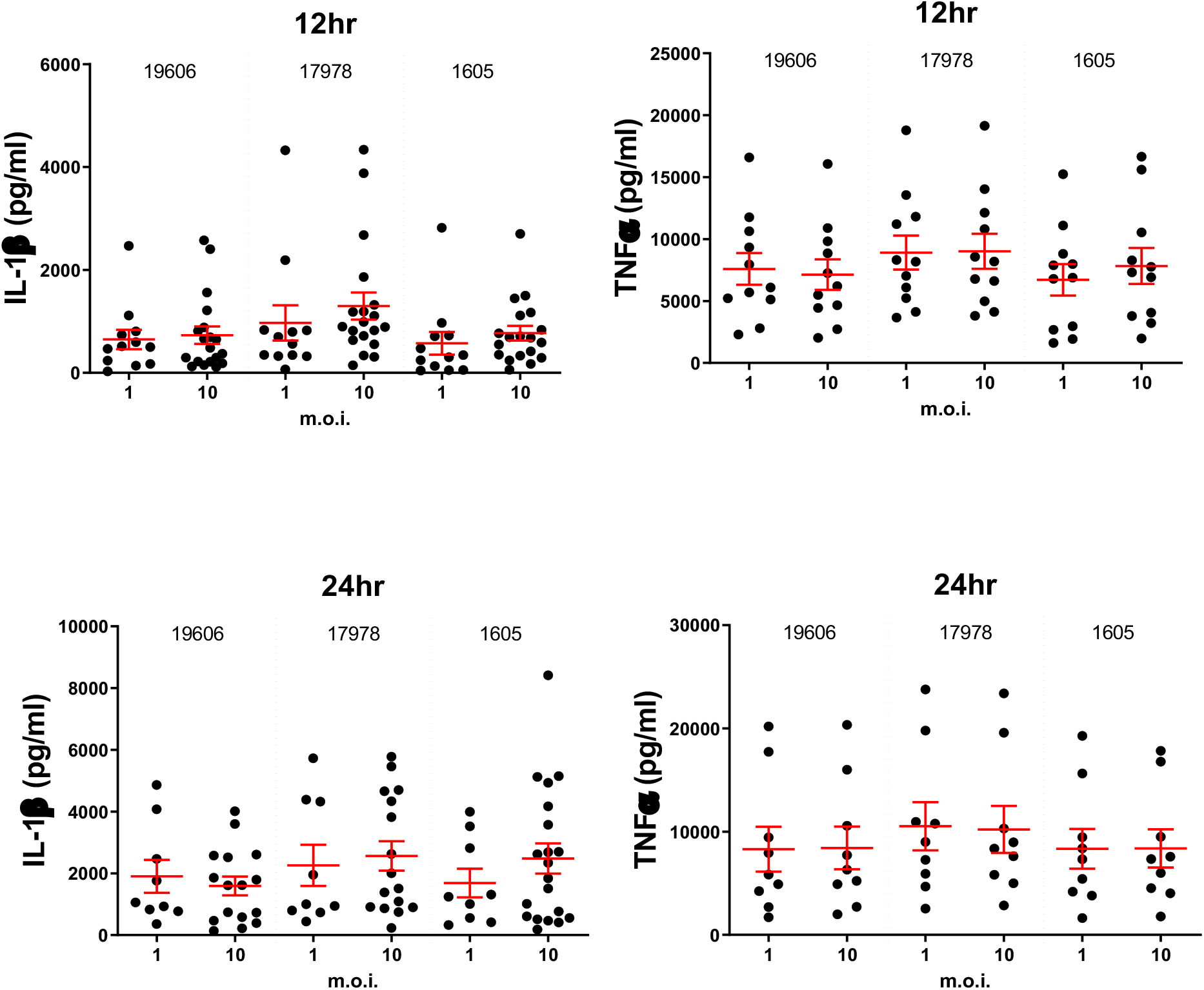
Mouse BMDMs were infected with indicated m.o.i. of live *A. baumannii* and analysed. ELISA of pro-inflammatory cytokines levels of TNFα and IL-1β produced by wild-type mouse BMDM after 12 and 24 hours of infection, n>3. mean ± SEM

**Supplementary Figure 2:**
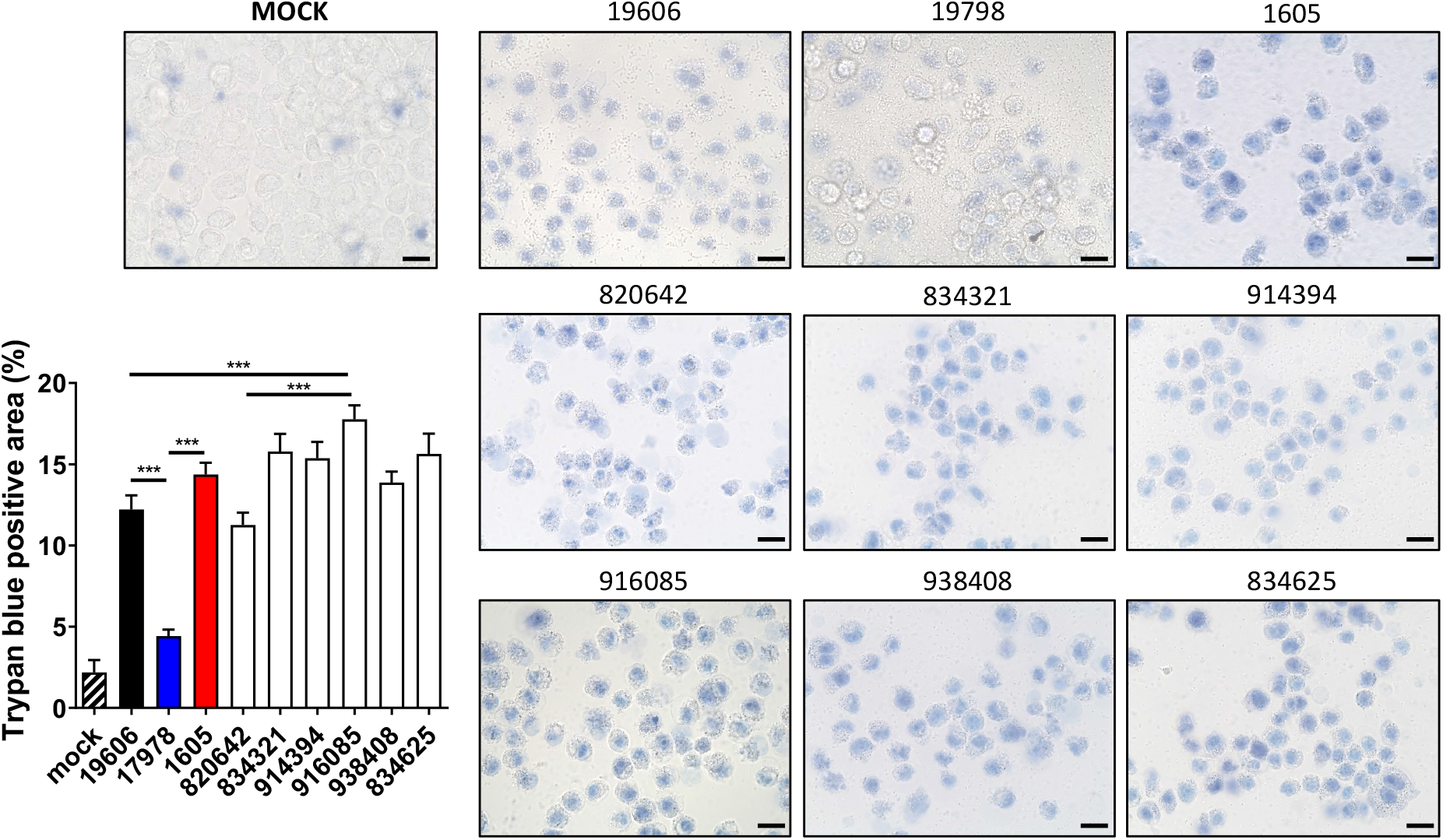
(a) Representative image and densitometry quantification of A549 cell death post *A. baumannii* infection at m.o.i. 80 for 24 hours, scale bar: 20 μm. n≥3. All data are shown as mean ± SEM. *, P < 0.05, **, P < 0.01, ***, P < 0.001 compared to wild-type unless indicated.

